# Microbial diversity of the Arabian Sea in the Oxygen minimum zones by metagenomics approach

**DOI:** 10.1101/731828

**Authors:** Mandar S. Paingankar, Kedar Ahire, Pawan Mishra, Shriram Rajpathak, Deepti D. Deobagkar

**Affiliations:** Molecular Biology Research Laboratory, Centre for Advanced Studies, Department of Zoology, Savitribai Phule Pune University, Pune - 411 007, India

**Keywords:** Arabian Sea, bacterial diversity, oxygen minimum zones, metagenomics, sulphur and nitrogen metabolism, Goa, Mangalore, Calicut

## Abstract

Large oxygen depleted areas known as oxygen minimum zones (OMZ) have been observed in the Arabian Sea and recent reports indicate that these areas are expanding at an alarming rate. In marine waters, oxygen depletion may also be related to global warming and the temperature rise, acidification and deoxygenation can lead to major consequences wherein the plants, fish and other biota will struggle to survive in the ecosystem.

The current study has identified the microbial community structure using NGS based metagenomics analysis in the water samples collected at different depth from the oxygen depleted and non-OMZ areas of Arabian Sea. Environmental variables such as depth, site of collection and oxygen concentration appeared to influence the species richness and evenness among microbial communities in these locations. Our observations clearly indicate that population dynamics of microbes consisting of nitrate reducers accompanied by sulphate reducers and sulphur oxidizers participate in the interconnected geochemical cycles of the OMZ areas. In addition to providing baseline data related to the diversity and microbial community dynamics in oxygen-depleted water in the OMZ; such analysis can provide insight into processes regulating productivity and ecological community structure of the ocean.

## INTRODUCTION

The Oxygen minimum zone (OMZ) in the Arabian-Sea is the second-most intense OMZ amongst the tropical oceans in the world^1,2^ with a near-total depletion of oxygen at depths from 200 to 1000m ^3^. In these locations, suboxic levels (≲5 μmol O2/kg) of oxygen are seen over vast areas at different depths and denitrification occurs in its upper portion^4^. Geochemical observations indicate that oxygen minimum zones have expanded over the past decades^5^ and could expand further in response to the ocean warming and increased stratification associated with climate change^6,7^. It has been suggested that the biological consumption of oxygen is most intense below the region of highest productivity in the western Arabian Sea^8–10^. The total volume of the OMZ in the ocean is growing at an alarming rate, their upper boundaries are vertically shoaling, and the degree of anoxia is intensifying within the cores of the OMZs^5,11^. The expansion of the oxygen minimum zones (OMZs) in the Arabian Sea has become the major concern because of its impact on the marine ecosystems. The expansion of the OMZs due to climate change and its impacts on the ecosystems and the atmosphere is multi-dimensional and requires intense study.

OMZ is characterized by high nitrite accumulations and very low or undetectable oxygen concentrations^12^. The nitrous oxide (N_2_O) concentration in the OMZ has been reported to vary inversely with nitrite concentration^13^. Often as the oxygen levels diminish the ecosystem cannot sustain normal biotic inhabitants and macrofauna. As a result, OMZs are often associated with coastal and equatorial upwelling regions and the increased primary production rates determine the high levels of altered microbial metabolism^11,14^. Importantly, Nitrogen (N) cycling plays crucial role in nitrate reduction to N2 (denitrification) and anaerobic ammonia oxidation (anammox) along with nitrate reduction to ammonia^15^. Moreover, nitrification has been shown to be an important source of oxidized N at the OMZ boundaries^16–18^.

Interestingly, various metagenomic studies on OMZ have revealed that complex communities (such as nitrifiers) play an important role in N cycle in the OMZ^18^. Members of the Planctomycetes, Thaumarchaeota and Nitrospinae phyla have been observed to perform the majority of anammox, ammonia oxidation and nitrite oxidation and play important role in the OMZ dynamics^12,18–24^. Although some reports exist, the denitrification^25–27^ and heterotrophic denitrification via a complete sequential reduction of nitrate (NO_3_) to N_2_ has not been fully explored in the OMZ areas^28,29^. A few studies have been carried out to understand the microbial diversity in the OMZ areas of Arabian Sea ^30–34^ The special growth requirements of these microbes and abundance of uncultured organisms (over 99%) make NGS based metagenomic the method of choice in order to unravel the complexities of microbial communities, their dynamics and ecological significance.

In the current study, water samples collected from different depths of sea (100 to 1000 meters across transect) from Goa, Mangalore and Calicut (Supplementary Information Table SI1) were processed for high throughput next generation sequencing based metagenomics (based on 16S rRNA gene sequencing). The microbial diversity and predicted metabolic activities associated with these microbial communities in OMZ and non-OMZ areas in Arabian Sea of India provide valuable insight into the nature of biogeochemical processes.

## MATERIAL AND METHODS

### Sample Collection

Water samples at different depths were collected during the Sagar Sampada cruise (Sagar Sampada Cruse Number 340, 16 May 2015 to 08 June 2015) from Goa (GAS1, GAS2, GAS3 and GAS4; distance from coast ranging from 51 km to 90 km), Mangalore (MGS5, MGS6, MGS7 and MGS8; distance from coast ranging from 52 km to 84 km) and Calicut (CLS9, CLS10 and CLS11; distance from coast ranging from 66 km to 109 km) (Supplementary Information Table SI1; Fig. 1) A conductivity–temperature–depth (CTD) system equipped with attached oxygen and turbidity measurement sensors was deployed to record the physical properties of the water (Supplementary Information Table SI2) and the samples were grouped into OMZ and non OMZ.

**Fig. 1.**
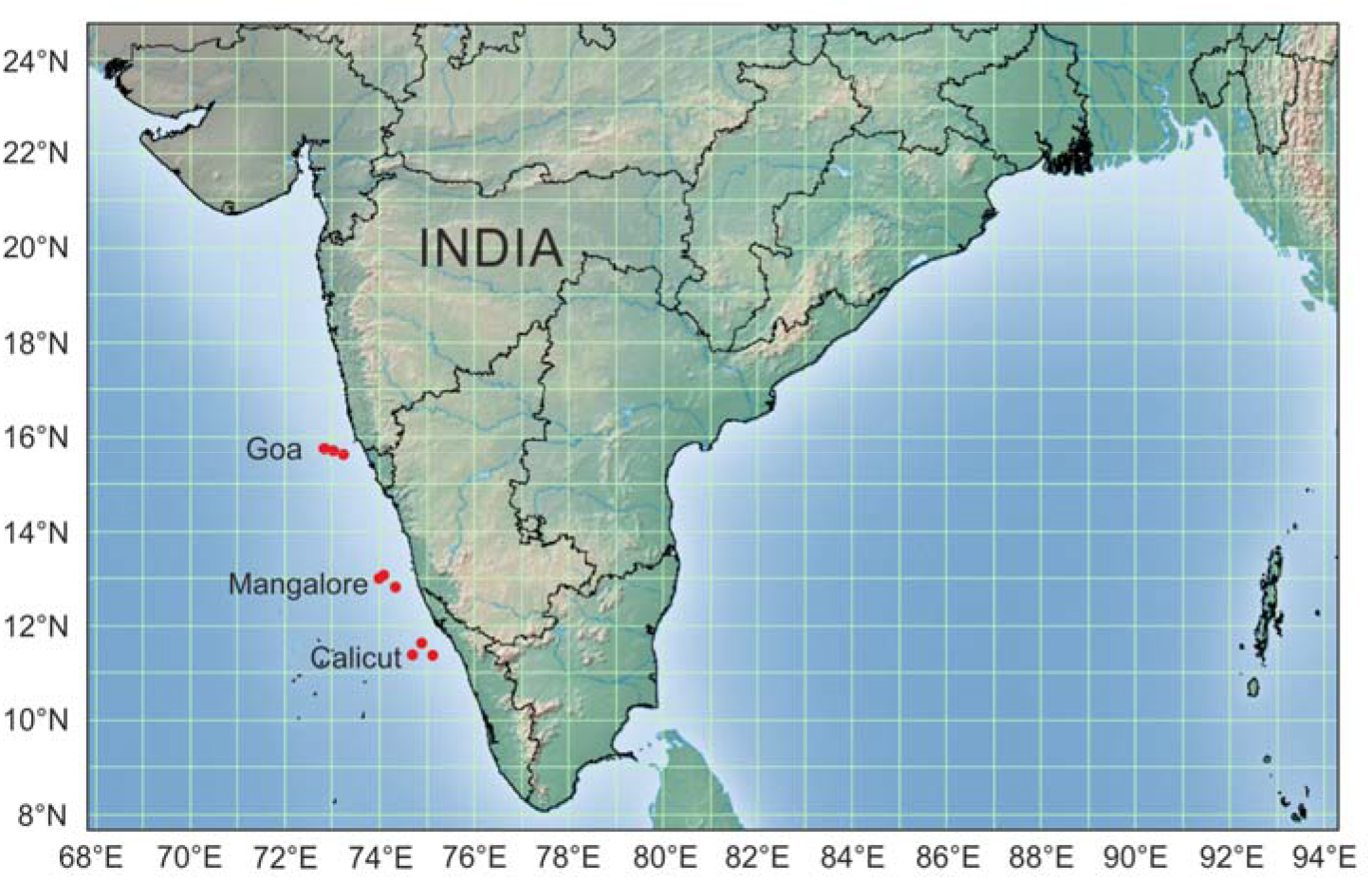
Location of sampling sites in Arabian Sea. (Distance from coast is depicted in Supplementary Information Table SI1)

### DNA extraction

1000 ml water was collected from each sampling sites and organisms collected by filtering water through 0.22 μm filter (Millex, Merck Millipore, USA) were utilised for DNA isolation using Power water DNA isolation kit (MoBio laboratories Inc. Carlsbad, CA). DNA isolation was carried out on the ship to avoid degradation of DNA. DNA concentration was measured using the Quantus fluorimeter (Promega, USA).

### Amplification primers and Sequence analysis

16s rRNA (corresponding to V3 and V4 regions) was amplified from total genomic DNA isolated (16S Amplicon PCR Forward Primer 5’TCGTCGGCAGCGTCAGATGTGTATAAGAGACAGCCTACGGGNGGCWGCAG3’; 16S Amplicon PCR Reverse Primer 5’GTCTCGTGGGCTCGGAGATGTGTATAAGAGACAGGACTACHVGGGTATCTAA TCC3’) with appropriate sample bar coding index sequences and Illumina adapters. AMpure XP beads were employed to remove unused primers and other unwanted nucleic acid fragments and thepurified PCR amplicons were quantified, normalized and an equimolar pool of all the samples was made. This multiplexed library was further subjected to QC using an Agilent Bioanalyzer DNA Chip. The sequencing libraries generated from V3 and V4 amplicons from all the samples were sequenced using an Illumina paired end overlapping sequencing. Sequence reads were binned according to index sequences and QC of the raw sequence data was performed by custom scripts. Low quality reads were filtered out and trimmed based on observed quality pattern in the data set. Read pairs with high sequence quality and overlapping regions were fused together to obtain a single read traversing full length of V3 and V4 region.

### Bioinformatics analysis

The sequences which were less than 300 bps and sequences with less than average quality score (25 or less) were removed from the library. The taxonomic assignment of unassembled clean metagenomic sequences was performed using Ez-Taxone database^35^ and BLASTX. Information related to the metagenomics reads of the samples is depicted in Table 1.

**TABLE 1.**
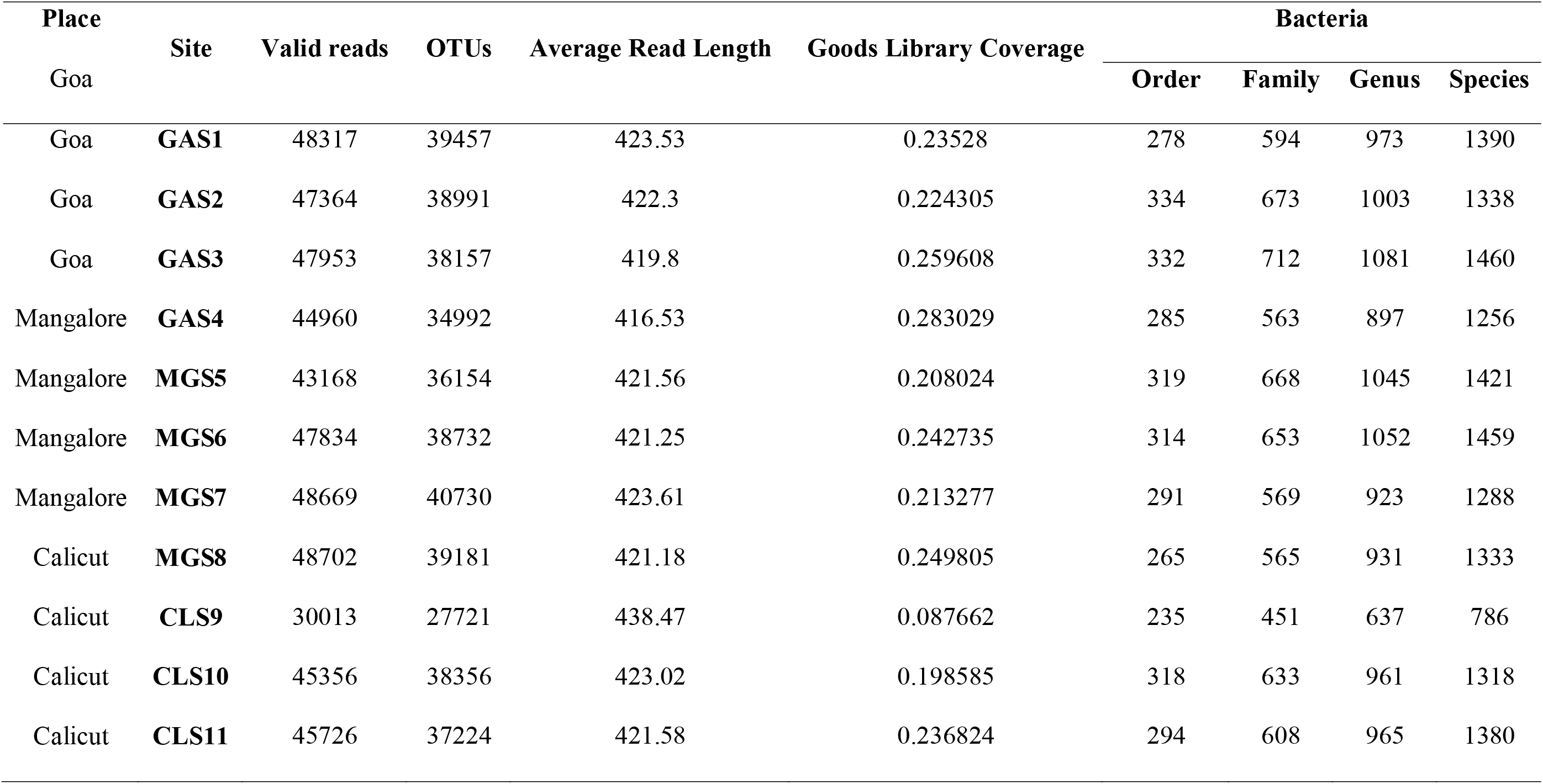
Metagenomics reads information and taxonomic affiliations of bacteria present in samples collected from Arabian Sea

| Place | Site | Valid reads | OTUs | Average Read Length | Goods Library Coverage | Bacteria |  |  |  |
| --- | --- | --- | --- | --- | --- | --- | --- | --- | --- |
|  |  |  |  |  |  | Order | Family | Genus | Species |
| Goa |  |  |  |  |  |  |  |  |  |
| Goa | **GAS1** | 48317 | 39457 | 423.53 | 0.23528 | 278 | 594 | 973 | 1390 |
| Goa | **GAS2** | 47364 | 38991 | 422.3 | 0.224305 | 334 | 673 | 1003 | 1338 |
| Goa | **GAS3** | 47953 | 38157 | 419.8 | 0.259608 | 332 | 712 | 1081 | 1460 |
| Mangalore | **GAS4** | 44960 | 34992 | 416.53 | 0.283029 | 285 | 563 | 897 | 1256 |
| Mangalore | **MGS5** | 43168 | 36154 | 421.56 | 0.208024 | 319 | 668 | 1045 | 1421 |
| Mangalore | **MGS6** | 47834 | 38732 | 421.25 | 0.242735 | 314 | 653 | 1052 | 1459 |
| Mangalore | **MGS7** | 48669 | 40730 | 423.61 | 0.213277 | 291 | 569 | 923 | 1288 |
| Calicut | **MGS8** | 48702 | 39181 | 421.18 | 0.249805 | 265 | 565 | 931 | 1333 |
| Calicut | **CLS9** | 30013 | 27721 | 438.47 | 0.087662 | 235 | 451 | 637 | 786 |
| Calicut | **CLS10** | 45356 | 38356 | 423.02 | 0.198585 | 318 | 633 | 961 | 1318 |
| Calicut | **CLS11** | 45726 | 37224 | 421.58 | 0.236824 | 294 | 608 | 965 | 1380 |

### Statistical analysis

Dominance, Simpson, Shannon, Evenness, Brillouin, Menhinick, Margalef, Equitability, Fisher_alpha, Berger-Parker, Chao-1, Whittaker, Harrison, Cody, Routledge, Wilson-Shmida, Mourelle, Harrison 2 and Williams indices of clonal and beta diversity were estimated using the PAST3 programs available from the University of Oslo website link Relationship between chemical composition and (i) species diversity unifrac distances, (ii) species alpha diversity indices and (iii) species Beta diversity indices were determined by Mantel tests. P values were calculated using 9999 permutations on rows and columns of dissimilarity matrices. Principal coordinate analysis (PCO), canonical correlation analysis (CCor), permutational analysis of variance (PERMANOVA) and analysis of similarity (ANOSIM) was performed using the Past 3 software. For the predictive functional analyses, the PICRUSt software package^36^ was used to identify predicted gene families and associated pathways.

### Analysis of predicated functional profiles for the identified microbial communities

The 16S rRNA sequencing data sets were analysed by PICRUSt script-(normalize_by_copy_number.py script) for copy number normalization^36^. Functional predictions were assigned up to KO tier 3 and categories including metabolism, genetic information processing, environmental information processing, and cellular processes were analysed further. KEGG Pathway analysis was carried out by employing functions .py PICRUSt scripts followed by STAMP (Statistical Analysis of Metagenomic Profiles) software^37^, with Welche’s t-test and P value cut-off of 0.05 was considered to reject null hypotheses. This identification of functional features of the genes and metabolic pathways has relevance in understanding metabolic processes in the context of the ecosystem.

### Identification of bacterial markers by LDA Effect Size (LEfSe) analysis

Linear Discriminant Analysis (LDA) Effect Size (LEfSe) analysis was utilised for identification of unique microbial communities present in different samples^38^. The LEfSe analysis with LDA score threshold of 2 using online Galaxy version 1.0 was used to identify variations in bacterial diversity at specific locations and depths.

## 3. RESULTS

### 3.1 Species diversity in Arabian Sea

Water samples (total 11) collected from Arabian Sea at specific locations and depths (Supplementary Information Table SI1, SI3) were subjected to metagenomic analysis using next generation sequencing technology of amplified rDNA libraries. A total of 498062 (45278.36 ±5369.15 per sample) high-quality sequences with 3551 (1311.72±186.45 per sample) distinct bacterial species were recorded (Supplementary Information Table 3) were identified. This data was analysed extensively to identify if there were differences in the OMZ and the non OMZ regions. In all three sampling sites in OMZ 1371 species were common. while 777 species were found common at all depths (100m, 200m, 500m and 1000m) across different sampling sites (Fig. 2A, 2B).

**Fig. 2.**
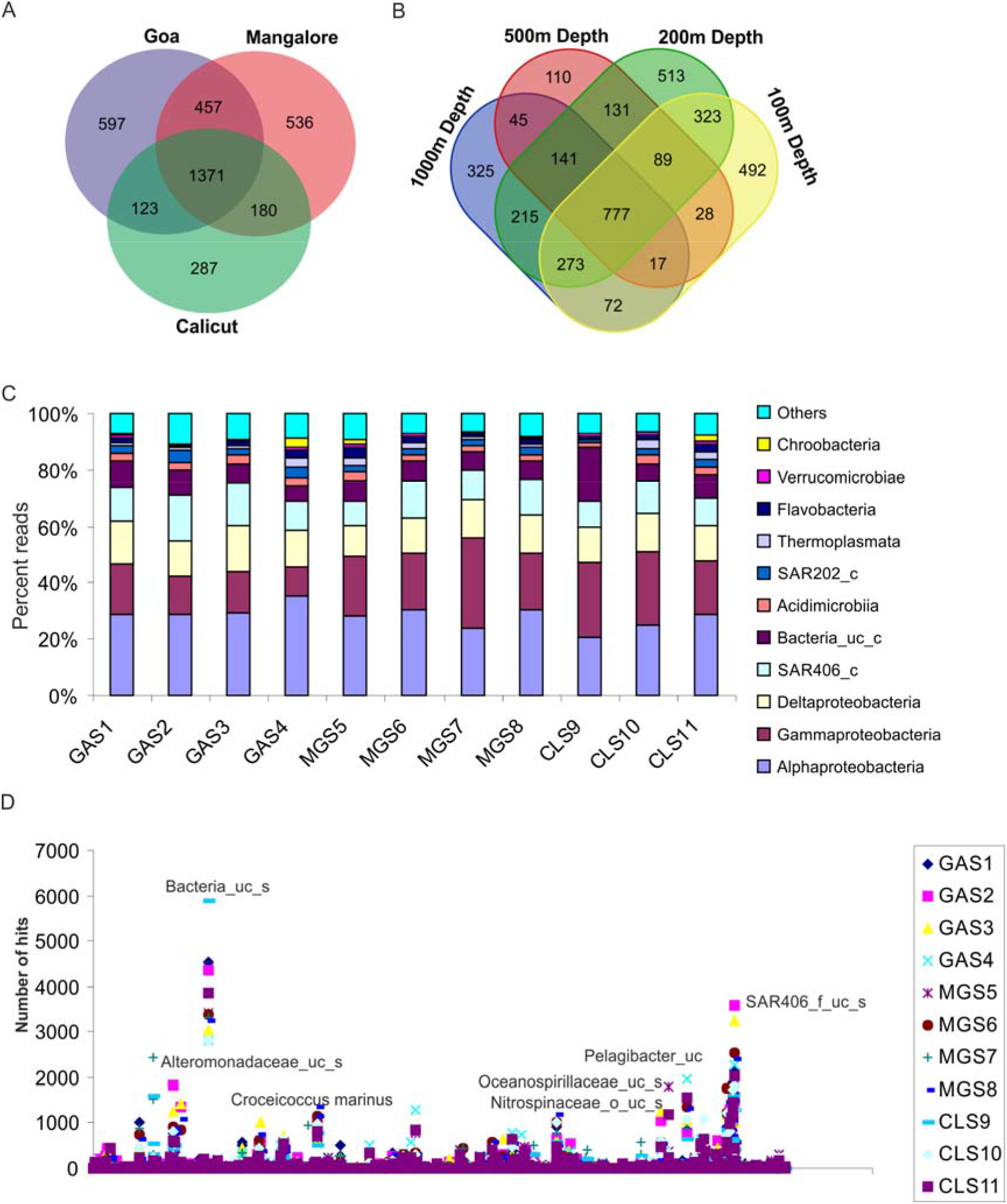
Venn diagrams showing the number of unique and shared species between the (**A**) three sampling sites (**B**) different depth of sea. Distribution of predominant bacterial class in sample based on 16S rRNA gene sequencing. (**C**) Phyla distribution. Observations are displayed as stacked bar charts for individual mangrove sample (x-axis) against the taxa abundance (y-axis). (**D**) Abundance of species. Observations are displayed as scatter plot for individual sample (x-axis) against the species abundance (y-axis).

#### Community composition of the Arabian Sea

A large number of uncultured and novel microbes were abundant at these locations (Supplementary Information SI3). Proteobacteria and SAR406 were common while Firmicutes, Spirochaetes, Chloroflexi and Verrucomicrobia were present in relatively lower numbers. Alpha proteobacteria (20.43-35.51%) (Fig. 2C) Deltaproteobacteria (11.03-15.93%) and Gamma proteobacteria (9.98-32.18%) were abundant in significant numbers in all the OMZ samples analysed. At the family level, SAR11-2_f (5.38-12.49%), Bacteria_uc_f (5.84-19.38%), Ruthia_f (0.69-7.56%), Arenicella_f (1.44-4.38%), Nitrospinaceae (1.86-5.89%), Erythrobacteraceae (0.45-4.80%) were present across all samples (Fig. 2B). Bacterial orders such as SAR11 (10.73-23.81%), Bacteria_uc_o (5.84-19.39%), Ruthia (0.70-7.91%), Alteromonadales (0.34-14.17%), Nitrospinaceae (1.96-6.11%) showed high abundance in all the samples. At 1000m depth SAR324_f (7.90-10.67%), Bacteria_uc_f (6.69-19.34%) and Erythrobacteraceae (0.94-4.80%) were predominant. Bacterial families such as Homogoneae and Thoreales were affiliated only with GAS4 sample whereas Synarophyceae, Ceramiales, Euglenida and Cloacamonas were exclusively present in CLS11 sample. Vaucheriales, Crenarchaeota, Pedinophyceae, Zetaproteobacteria and synergista were specific to MGS5 sample. Nitrospireae, Methanomicrobia, Bryospida were exclusive to 200m depth. SAR11-2_f (7.81-12.30%) and SAR11-1_f (6.31-12.32%) were predominant at 100m and 200m depth while Prochlorococcaceae (1.81-3.19%) was predominantly present at 100m depth. SAR406_o_uc (1.17-2.69%) was abundant at 200m depth.

Genera such as *Bacteria_uc_g* (5.85-19.38%), *Pelagibacter* (3.44-9.89%), *SAR324_g* (2.60-7.20%) were ubiquitous. *Croceicoccus* (1.25-2.14%) was predominantly present in samples from Goa as compared to other samples. Correspondence analysis revealed that in the depth of 1000 meters, *Methylopila, Mycoplasma, Asticcacaulis, Cellulomonas, Phalanopsis* were exclusive to MGS7 sample whereas *Spirochaeta, Chroococcidiopsis, Thysira, Leeuwenhoekiella* were selectively present in CLS9 sample. Water sample at 1000 meters depth from Mangalore (MGS5) revealed presence of *Terasakiella, Chlorodendrales, Vaucheria, Congregibacter, Planktotelea, Pseudoflavinifactor* whereas *Spirobacillus, Moraxella, Tiobacter, Roseburia, Marinoscillum, Thiohalophilus, Akkermansia, Caedobacter, Oceanicaulis, Epibacterium, Ditylium* were exclusively seen in Goa (GAS4). A more detailed analysis of data based at the species level revealed that *Bacteria_uc_s, SAR406_f_uc_s, Ruthia_f_uc_s, Arenicella_f_uc_s, Nitrospinaceae_uc_s, Oceanospirillaceae_uc_s* and *Rhodospirillaceae_uc_s* were present in high numbers in all samples that were analysed in our study (Fig. 2D, Supplementary Information Table SI3).

#### Linear Discriminant Analysis (LDA) Effect Size (LEfSe) analysis

In order to determine the unique and predominant bacteria present at a particular location, a comparative assessment of the biodiversity LefSe was carried out. This resulted in the identification of specific marker families for different locations as well as for the various sampling depths (Supplementary information SI2; SI3). Bacterial families including Erysipelotrichi_uc_f, SAR11_uc, EU335161_o_uc, Pseudoalteromonadaceae and Alteromonadales_uc were specific to Calicut while family FJ444691_c_uc_f was seen in Mangalore. Water samples from Goa Salinisphaeraceae, EU686587_f, and Dehalococcoidales_uc showed significant enrichment. LefSe analysis with respect to the depth showed enrichment of bacterial families (total 66) such as Brumimicrobiaceae, Bacteriovoracaceae, Dinophysiaceae, Spirochaetaceae, and Chaetocerotaceae at shallower depth (100m) while bacterial families (In total 22) such as Methylobacteriaceae, Halomonadaceae and Rhizobiaceae were found to be enriched at the depth of 500-1000m.

#### Alpha and Beta diversity of samples

Alpha diversity analysis highlighted the rich taxonomic diversity in the sea samples (Supplementary Information Table SI4). Simpson index of all samples close to 1 for all samples indicated the presence of highly diverse microbial communities in samples. Shannon’s index varied from 4.29 to 5.21 indicating the high species richness in bacterial diversity in these sea samples. Evenness index ranged from 0.093 to 0.138 while Margalef richness index was also high emphasising the richness of bacterial species in the sea area. Chao-1 analysis predicted the number of bacterial species in each sample to be between 1106-2122 (Supplementary Information Table SI4). No significant difference was observed in alpha diversity indices when pair wise comparison was carried out between Goa, Mangalore and Calicut sampling sites (ANOVA P>0.05; Mann Whitney U test P>0.05 for each comparison). Beta diversity indices of these sea samples are depicted in Supplementary Information Table SI5. At species level, high beta diversity was observed in all the sea samples (Supplementary Information Table SI5). This extensive analysis documented not only the rich and diverse micro flora present in each sample but also emphasised the differences in the microbial communities in the Arabian Sea.

#### Depth and geochemical parameter influences the community structure

Principal component analysis (PCoA) led to the identification of depth as an important determinant which influences characteristic and typical community structures of a given niche. (Fig. 3) Samples from similar depth clustered together, indicating that the communities in these locations are very similar to each other. The correlations between environmental factors and alpha diversity indices were accessed by Canonical correlation analysis (CCorA) (Fig. 4A). Depth, turbidity and density were seen to influence the dominance of certain species while temperature and conductivity correlated with the richness and evenness in samples (Fig. 4 A). Beta diversity showed a correlation with geochemical characters of sample (r = −0.262; p value = 0.05) while alpha diversity (r = −0.0004; p value = 0.744) and unifrac distances among the sampling sites (r = −0.09; p = 0.517) were not affected by the geochemical characters of sample (Fig. 4B). The sampling site did not influence the community composition while depth was a major factor (PERMANOVA (F= 4.036 *P*=0.0009ANOSIM R= 0.7222, *P*= 0.0008) (Table 2).

**Fig. 3.**
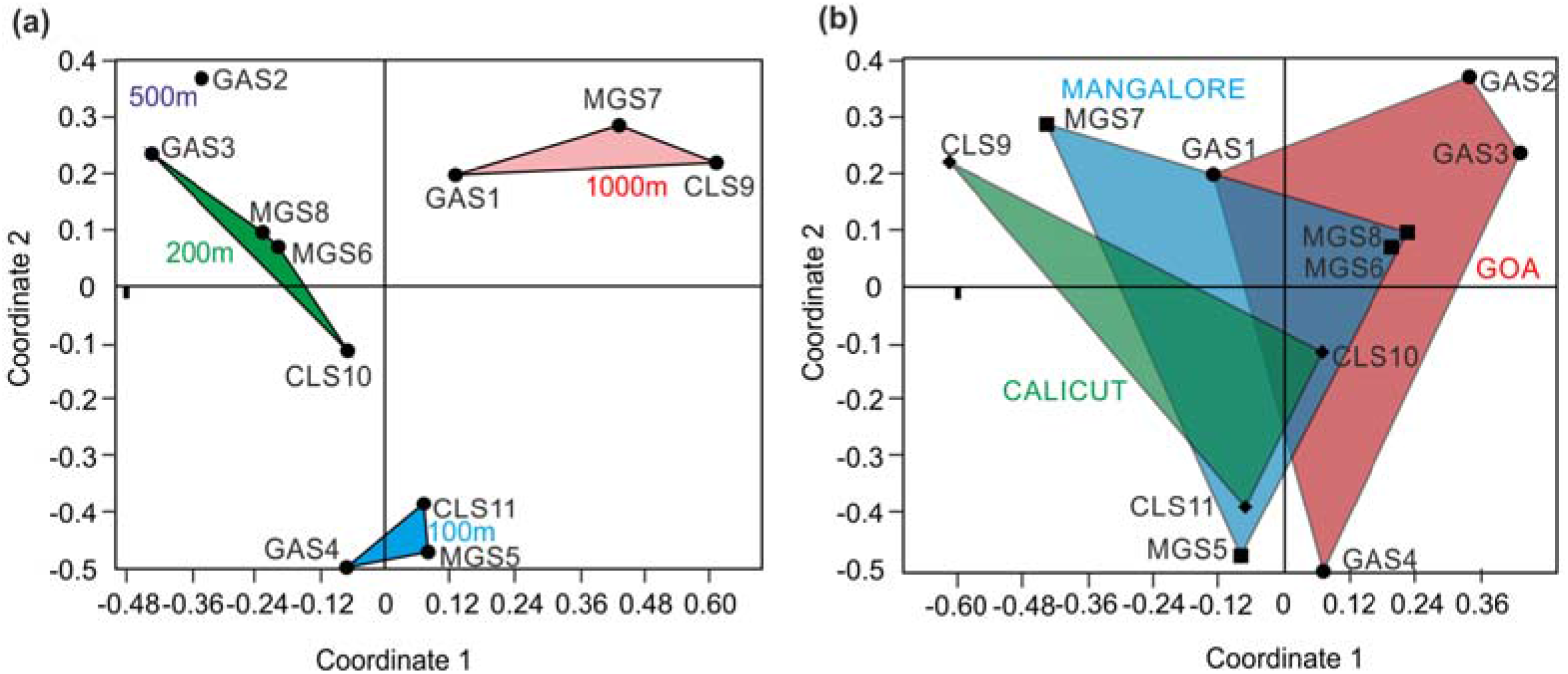
Principal Coordinates Analysis (PCoA) representation of the similarity matrix generated by cluster analysis. Samples from (**A**) depth and (**B**) collection site are represented by a different shape, and the distance between dots represents relatedness obtained from the similarity matrix.

**Fig. 4.**
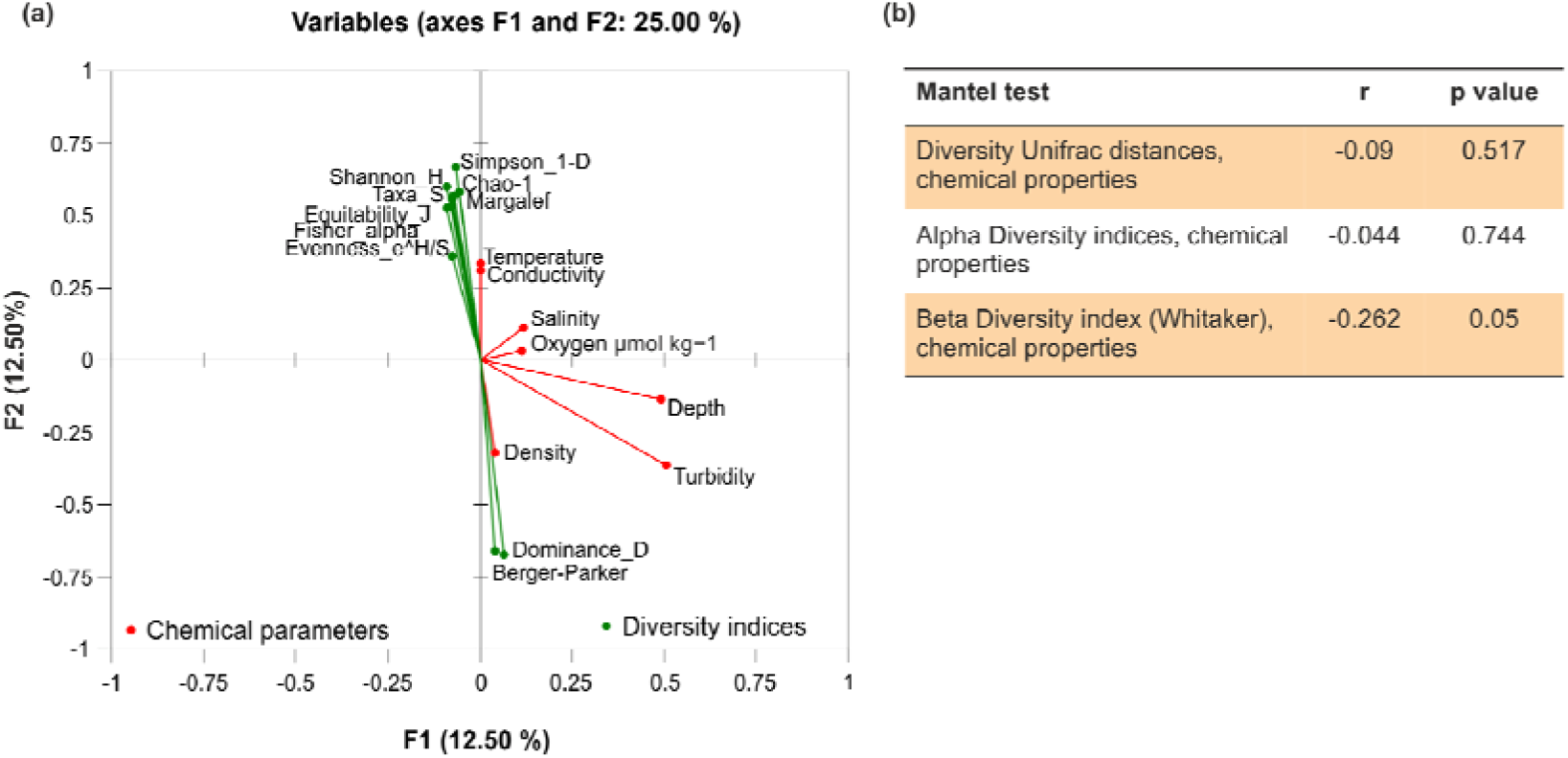
Association of environmental parameters and diversity indices (A) Canonical correlation analysis (CCorA) (B) Mantel Test

**TABLE 2.** Effect of depth and sampling location on bacterial diversity of samples collected from Arabian Sea

| Test | F | P |
| --- | --- | --- |
| **Permanova** |  |  |
| Sampling Site | 1.041 | 0.416 |
| Depth | **3.503** | **0.0002** |
| **Pair wise comparison (t test)** |  |  |
| Goa vs Magalore | 0.9932 | 0.5742 |
| Goa vs Calicut | 1.294 | 0.2914 |
| Calicut vs Mangalore | 0.8273 | 0.6308 |
| 100m vs 200m | **4.952** | **0.0278** |
| 100m vs 500m | 5.166 | 0.2496 |
| 100m vs 1000m | 3.771 | 0.1022 |
| 200m vs 500m | 2.472 | 0.1949 |
| 200m vs 1000m | **3.668** | **0.0273** |
| 500m vs 1000m | 1.75 | 0.2585 |

#### OMZ vs Non-OMZ samples

A comparative analysis and assessment of all samples showed presence of 2718 species in OMZ areas and 2223 species in non OMZ. It was seen that 1690 operational taxonomic units (OTUs) were common in OMZ and non OMZ while 1328 and 533 are unique to OMZ and non-OMZ respectively (Fig. 5A). This clearly documents that although several common inhabitants are seen in the ocean the depletion of oxygen is changing the species pattern. Differential abundance was clearly visible when the top 50 families present at these locations were compared (Supplementary Information Table SI3). In case of OMZ, SAR324, Ruthia, Arenicella, Zunongwangia, Rhodospirallacae, Nirtitreductor, Oleiobacter, Hyphomonas, Methylophaga, Clamydiales, Xanthomonadeacae, Sphingopyaix, Pararhodobacter, Anoxybacilllus, Gemella, Phenylobaterium and Legionella families were present while in non-OMZ Prochorobacteriacae, SAR11, Dianophyaceae, pelagomonadeacae, Delatproteobacteria, Firmicutes, Cytophagales, Flavobacteriales, Chroococcales, Dongicola, Planctomycetacia, Sphingobacteria, Draconibacterium familes were unique. Out of total 203 families which showed differential abundance, 86 were present in OMZ whereas 117 in Non-OMZ respectively (Fig. 5B). The principal component analysis (PCoA) revealed that non OMZ samples from similar depths clustered closely together; indicating that the communities in these locations are similar to each other and the depth typically influences characteristic and typical community structures.

**Fig. 5.**
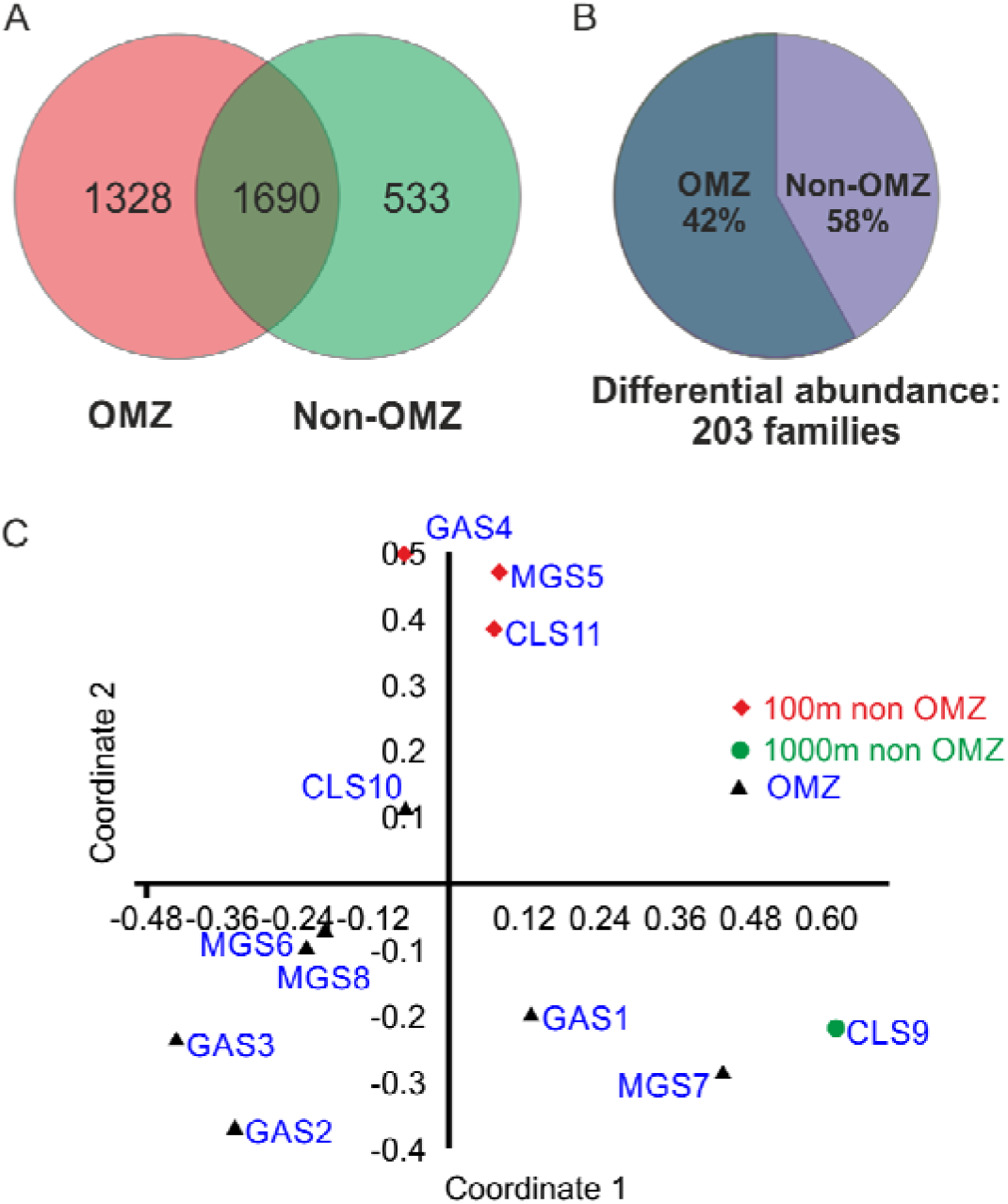
Comparison of differentially present microbial diversity in OMZ and non-OMZ areas

#### Functions associated with microbial communities

PICRUSt analysis is a bioinformatics software package designed to predict functional content from microbial community identification carried out by 16S rRNA based metagenomic analysis. The percent OTUs associated with different metabolic functions are xenobiotics biodegradation (2-3%), glycan biosynthesis (2.3-3 %), energy metabolism (7-7.5%) and (3-45%) for lipid metabolism. Three terms under Environmental Information Processing contain membrane transport (9-13%), signal transduction (1-2%) and signalling molecules and interaction (0.04-0.06%). DNA sequences encoding proteins such as Nitrogenase FeMo-cofactor scaffold and assembly protein NifN, QscR quorum-sensing control repressor, Cobalt-zinc-cadmium resistance protein CzcA; Cation efflux system protein CusA, tellurite resistance protein-related protein, Nitrite reductase [NAD(P)H] large subunit (EC 1.7.1.4), Nitrogenase FeMo-cofactor synthesis FeS core scaffold, Sulfur deprivation response regulator proteins and assembly protein NifB were found predominantly in samples from Goa as compared to other samples. In sea water samples from Mangalore, gene sequences encoding activities such as Cobalt-zinc-cadmium resistance protein CzcD, Sirohydrochlorin cobaltochelatase CbiK (EC 4.99.1.3) / Sirohydrochlorin ferrochelatase (EC 4.99.1.4), type IV fimbrial biogenesis protein PilY1, predicted L-rhamnose ABC transporter, methyl-accepting chemotaxis protein III, phage tail protein were enriched as compared to other activities. Predominance of genes for 5-O-(4-coumaroyl)-D-quinate/shikimate 3’-hydroxylase (EC 1.14.13.36), beta-glucanase precursor (EC 3.2.1.73) (Endo-beta-1,3-1,4 glucanase) (1,3-1,4-beta-D-glucan 4-glucanohydrolase), glutathione S-transferase C terminus, Shikimate/quinate 5-dehydrogenase I beta (EC 1.1.1.282), Tlr0729 protein, two component system response regulator, putative Fe-S containing oxidoreductase, possible polygalacturonase (EC 3.2.1.15), microbial collagenase (EC 3.4.24.3), chitosanase, aspartate ammonia-lyase (EC 4.3.1.1), FtsK/SpoIIIE family protein, putative EssC component of type VII secretion system were dominant in microbial communities from Goa and Calicut.

PICRUSt and STAMP analysis have identified OTUs associated with few of these KO terms differ significantly (P<0.05) between samples collected from different locations. On comparison of samples from Goa and Mangalore, Dioxin degradation and translation proteins differed significantly, while processes related to polycyclic aromatic hydrocarbon degradation enriched differentially in Calicut and Mangalore samples. In Calicut and Goa samples bacterial OTUs associated with Phenylpropanoid biosynthesis, polycyclic aromatic hydrocarbon degradation, Dioxin degradation showed differential enrichment.

## DISCUSSION

Arabian Sea is typically characterised by presence of vast areas of OMZs and these are expanding further. Depletion of oxygen in the habitats changes the microbial composition and leads to alterations in the nutrient as well as elemental cycles. Analysis of the correlation between the geochemical parameters and bacterial diversity is important in the understanding the dynamics of microbial communities in the OMZs. This study emphasises that Arabian Sea has high species richness with a complex community structure across oxygen gradients and between the depths of sea (Fig. 2). Chao-1 analysis highlighted presence of diverse assemblage of indigenous microbial species that remain completely uncharacterized at present. Our analysis indicated that relationships between environmental variables, conductivity, temperature and oxygen concentration have significant role in increasing the species richness and evenness in microbial communities (Fig. 4). OMZ samples in the Arabian sea displayed rich taxonomic diversity which typically showed a depth specific variation.

Nitrate reducing bacteria were present at all collection sites in OMZ and non OMZ areas of Arabian Sea. Reports from the suboxic zone of the Black Sea have identified single clade of nitrifying Crenarchaeota which is closely related to *Nitrosopumilus maritimus^39^.* Global Ocean Sampling (GOS) database across diverse physiochemical habitats and geographic locations has 1.2% *N. Maritimus*^40^. Interestingly, *N. Maritimus* is a cultured nitrifier isolated from a marine aquarium^41^. It has been shown that *N. maritimus* typically dominates low depth samples^21^. However, in the current study, *N. maritimus* was underrepresented in low depth samples.

Based on the metagenomic profiles of microbial assemblage gene repertoire and predicted functions were assessed. Genes encoding nitrite/nitrate sensor proteins, nitrilase, nitrate reductase, nitrate reductase associated proteins werepredominant in the datasets, emphasising that nitrate/nitrite metabolism plays a key role in the dynamics of microbial communities in the OMZ areas and play important role in nitrogen cycle in OMZ (Fig. 6). Naqvi *et al.* ^42^ have reported the presence of the nepheloid layer with significant amounts of suspended matter caused by bacteria in Arabian Sea while an increase in nitrifying bacteria (both ammonium and nitrite oxidizers) has been suggested to be the cause for such nepheloid layer^43^. Recent taxonomic, metagenomic, and metatranscriptomic analysis of many OMZs has shown that diverse sulphur-oxidizing microbial community are abundant and these communities are particularly enriched in *γ*-proteobacteria. Interestingly, the sulphate reducing bacteria (SRB) were present through-out the water column at all collection sites in our analysis. The presence of SRB has been shown not only at the bottom sediments but also in aerobic surface waters and beach sediments^44^. It has been shown that SRB populations increase from the surface waters up to the oxic–anoxic boundary. Colourless sulphur-oxidizing bacteria have earlier been reported from the Arabian Sea and these bacteria are known to mediate nitrogen cycle reductively even under autotrophic conditions^45^. SRBs are also known to participate in nitrate reduction. Jayakumar *et al*.^46^ and Ward *et al*.^47^ reported the dominance of denitrifying bacteria in the biomass of the OMZ and suggested that the denitrifying bacteria in this zone could be in a viable but non culturable state.

**Fig. 6.**
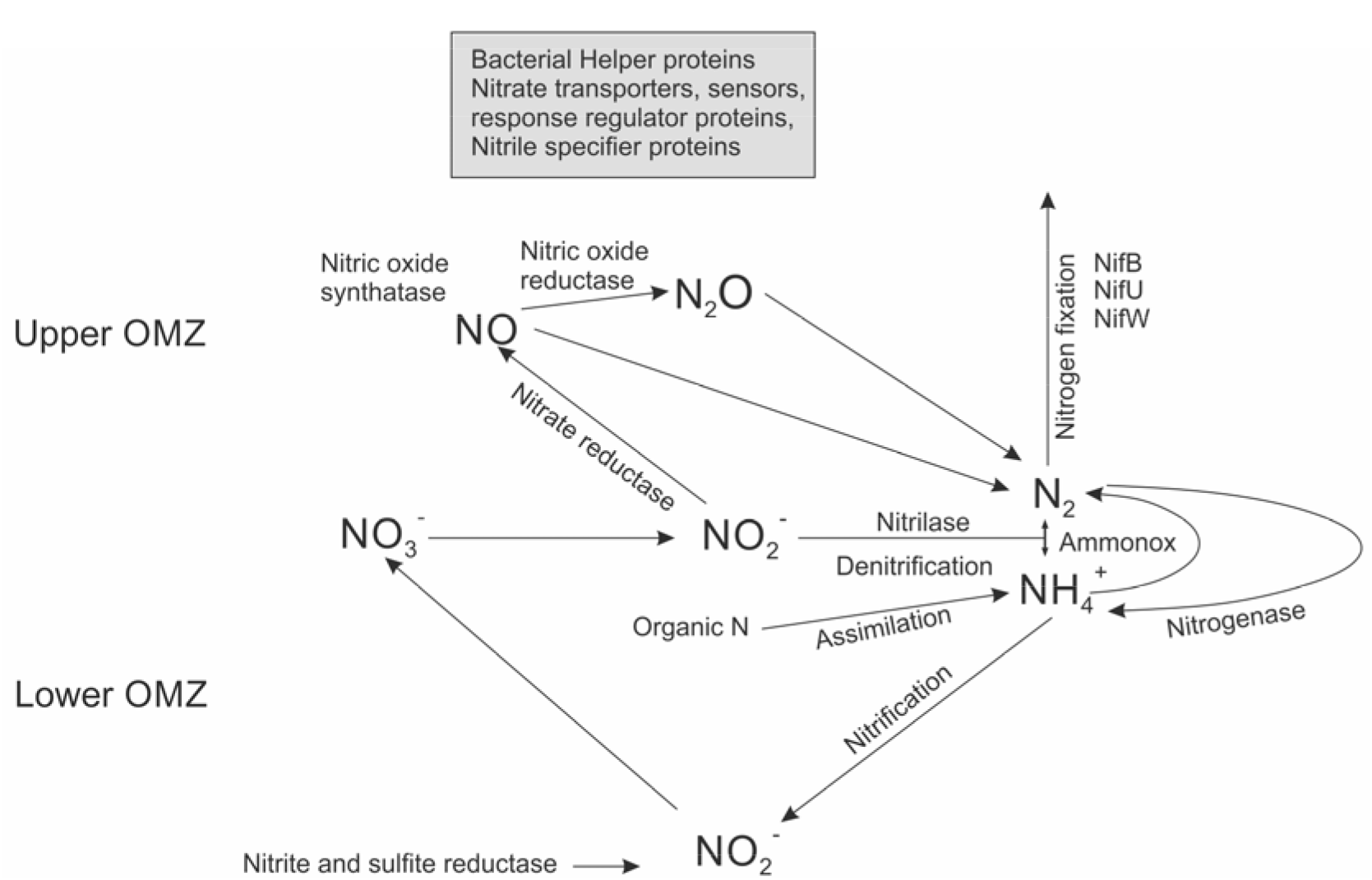
Probable nitrogen cycle in oxygen depleted zones in Arabian Sea.

Our analysis revealed that microorganisms involved in activities associated with sulphate metabolism were predominant with Sulphate permeases and reductase being predominant in the OMZ areas of Arabian Sea. Additionally, together with other recent analyses^48–50^, our results indicate the presence of an active sulphur oxidizing community in the Arabian sea OMZ. It is likely that sulphur cycle carried out by these SRB fuels nitrate reduction, thereby supplying additional substrates (nitrite and ammonia) for anammox bacteria. Comparative analysis of OMZ and Non-OMZ samples revealed that species such as *Zunongwangia profunda, Roseovarius nubinhibens, hydrocarbonoclasticus, Prochlorococcus marinus* and *Ruegeria pomeroyi* were common in oxygen depleted waters. These bacteria are known to be actively involved in dimethylsulfoniopropionate (DMSP) metabolism. This observation is important in the perspective of global climate change since DMS is thought to play a key role by decreasing the absorption of solar radiation and thereby influence temperature changes.

PICRUST analysis results obtained in the current study indicated presence of microorganisms harboring genes such as alkB, AlmA, CYP153A and AlkB are enriched in the OMZ in Arabian sea similar to the reports from Atlantic Ocean and Bay of Bengal ^51–53^. Presence of hydrocarbon degrading organisms points to presence of alkanes and hydrocarbon. In comparison to Calicut, Goa and Mangalore samples showed enrichment for polycyclic aromatic hydrocarbon degradation pathway suggesting anthropogenic activities in these areas. Deep sea water samples from depth 100-500 m depth were found to be enriched in (In total 22) Methylobacteriaceae, Halomonadaceae, Alcanivoracaceae and Rhizobiaceae. Methylobacterium which derive energy from the oxidation of thiosulfate to sulfate. The current study provides baseline data related to the diversity and potential microbial communities in oxygen-depleted water which could provide a basis for better understanding of the microbiological function, dynamics, and distribution in the oceanic OMZ. Further the results obtained in this study indicate the location specific functional divergence in bacterial community and therefore it would be interesting to carry out detailed functional analysis for bacterial diversity form Arabian Sea at more locations and with multiple samples in different seasons. Although our understanding of the OMZ and interplay between geochemical processes and microbes has been improving in recent years, the potential impacts of OMZs on marine ecosystem structure and global geochemical cycling remains to be completely elucidated. In this context we need to accelerate the exploration and discovery of microbes and their interplay with geochemical processes in the OMZ.

## Acknowledgement

We thank Prof. Dileep N. Deobagkar for valuable suggestions. The authors are grateful to Dr. M. Sudhakar, Dr. Saravannane, and Dr. R. A. Shivaji for their help during the sample collection.

## Author Contributions

DDD, MSP designed the experiments; DDD, KA, MSP, SR, PM carried out the work; DDD, MSP, SR, PM interpreted the results; DDD, MSP wrote the manuscript.

